# Addictions may be driven by competition-induced microbiome dysbiosis

**DOI:** 10.1101/2022.08.02.502262

**Authors:** Ohad Lewin-Epstein, Yanabah Jaques, Marcus W Feldman, Daniela Kaufer, Lilach Hadany

## Abstract

Recent studies revealed mechanisms by which the microbiome affects its host’s brain, behavior and wellbeing, and that dysbiosis – persistent microbiome-imbalance – is associated with the onset and progress of various chronic diseases, including addictive behaviors. Yet, understanding of the ecological and evolutionary processes that shape the host-microbiome ecosystem and affect the host state, is still limited. Here we propose that competition dynamics within the microbiome, associated with host-microbiome mutual regulation, may promote dysbiosis and aggravate addictive behaviors. We constructed a mathematical framework, modeling the dynamics of the host-microbiome ecosystem in response to alterations. We find that when this ecosystem is exposed to substantial perturbations, the microbiome may shift towards a composition that reinforces the new host state. Such positive feedback loop augments post-perturbation imbalances, hindering attempts to return to the initial equilibrium, thus promoting relapse episodes and prolonged addictions. We also find that the initial microbiome composition is a key factor: a diverse microbiome enhances the ecosystem’s resilience, whereas lower microbiome diversity is more prone to reach dysbiosis, exacerbating addictions. This framework provides novel evolutionary and ecological perspectives on host-microbiome interactions and their implications for host behavior and health, while offering verifiable predictions with potential relevance to clinical treatments.

## Introduction

Addiction is a brain disease where a victim experiences an uncontrollable motivation to engage in a rewarding behavior despite the behavior’s harmful consequences ^1^. Addiction encompasses a range of substance abuse disorders including various drugs, alcohol and cigarettes, as well as excessive food consumption. In recent years studies have found various associations between host addictive behavior, and the composition and functioning of the microbiome, the collection of microorganisms residing in the host ^2-6^. These studies accord with a vast body of microbiome research that has revealed pathways by which the microbiome can affect its host’s health and behavior, and others through which the host shapes the microbiome community. Despite these findings, understanding of the ecological and evolutionary processes that shape the host-microbiome ecosystem, and the extent of the microbiome involvement in host chronic diseases and addictive behaviors is still limited.

We hypothesize that microbial strains that are part of the host microbiome, may have evolved to affect the host state in a way that improves their condition within the microbiome community. This may lead to a community of microbes that affect the host in different directions. Host addictive behavior produces an alteration in the environmental conditions of its resident microbes. Even if the addiction is largely deleterious to the microbes (e.g., via toxins), for some microbes it may be less deleterious than to others, therefore generating a shift in the microbial selection regime, and perturbing the microbiome composition ^7-11^. Strains that proliferate in the new conditions might benefit from encouraging the host to continue its new behavior. Thus, the microbiome may play a significant role in enhancing and maintaining addictive behavior.

Addiction is characterized by both negative and positive emotional states attributable, at different stages, to alterations in activity of neurotransmitters: a binge/intoxication stage in which the mesolimbic dopamine system, a key part in the reward circuitry, produces reinforcing actions; a withdrawal stage which is associated with alterations in neurotransmission in the amygdala that generate emotional stress; a preoccupation/anticipation stage in which dysregulation of prefrontal cortex and insula projections interrupts control over incentive salience and therefore goal-directed behavior. The microbiome has been shown to affect the host brain in multiple ways, for example by modulation of neurotransmitters and interaction with the central nervous system via the gut-brain axis ^12-15^. Through this modulation, the microbiome can influence neural activities that are involved in brain reward and withdrawal circuitry by generating negative and positive feedback loops, thus promoting addictive behaviors. For example, neurotransmitters crucial to the functioning of these circuits, such as dopamine, GABA, and serotonin were found to be produced or regulated by the gut microbiome ^16-20^.

On the other hand, consumption of various addictive substances, including consumption of opioids ^21,22^, alcohol ^7,8^, smoking ^9,23^ as well as certain diets ^24,25^, has been linked to alterations in microbiome composition. Every such alteration involves a decrease in the frequency of some microbes, and an increase in the frequency of others. From the perspective of the latter, the host behavior causing the alteration is a beneficial one.

Previous models studied interactions between microbial strains within the microbiome ^26-28^ and also host-microbe interactions ^29-32^. Here we combine these two approaches into a framework that models the eco-system of a host, whose wellbeing and behavior alters the microbiome composition, and the microbiome, which affects the host state. Using this model we analyze several aspects of addictive behavior, focusing on the potential effects of the microbiome on addiction initiation and withdrawal. We show that within-microbiome competition may drive the evolution of microbial feedback on host behavior in a way that improves the microbe’s competitive state but may also be involved in exacerbating addictive behaviors. Our model also suggests that microbiome richness and functional diversity are key factors affecting the likelihood of reaching dysbiosis. We find that reduced microbiome richness and diversity facilitate a less resilient host-microbiome ecosystem, which is more likely to destabilize and reach dysbiosis, hence aggravating and prolonging the process of behavioral withdrawal and increasing the frequency of relapse episodes.

## Results

We suppose that microbes can secrete products that affect functions of the host reward and withdrawal circuitry, generating either positive or negative feedbacks, which affect the trajectory of the host’s behavior. We first examine the evolution of microbes that can affect their host’s behavior. For this purpose, we model the growth dynamics of two microbial strains that reside in a host and compete over resources: one strain affects its host behavior, and the second does not. We then extend the model to account for competitive dynamics among numerous microbial strains that can affect host behavior. We investigate the host-microbiome interactions, focusing the analysis on host addictive behavior (see Methods for model description).

We find that a microbe that affects feedback to host behavior may be selectively favored over a wide parameter range, including when it suffers a cost for producing this feedback. Such microbial effects would be favored when the additional resources the microbe gains from the altered host behavior, either directly or through the competition with other strains, compensate for the costs required to produce the effects. Moreover, once the affecting strain manages to draw its host towards a behavior that is more beneficial to that strain, its proportion in the microbiome rises and so does its ability to continue affecting the host, further inducing the host to continue the new behavior (Fig. 1).m

**Figure 1.**
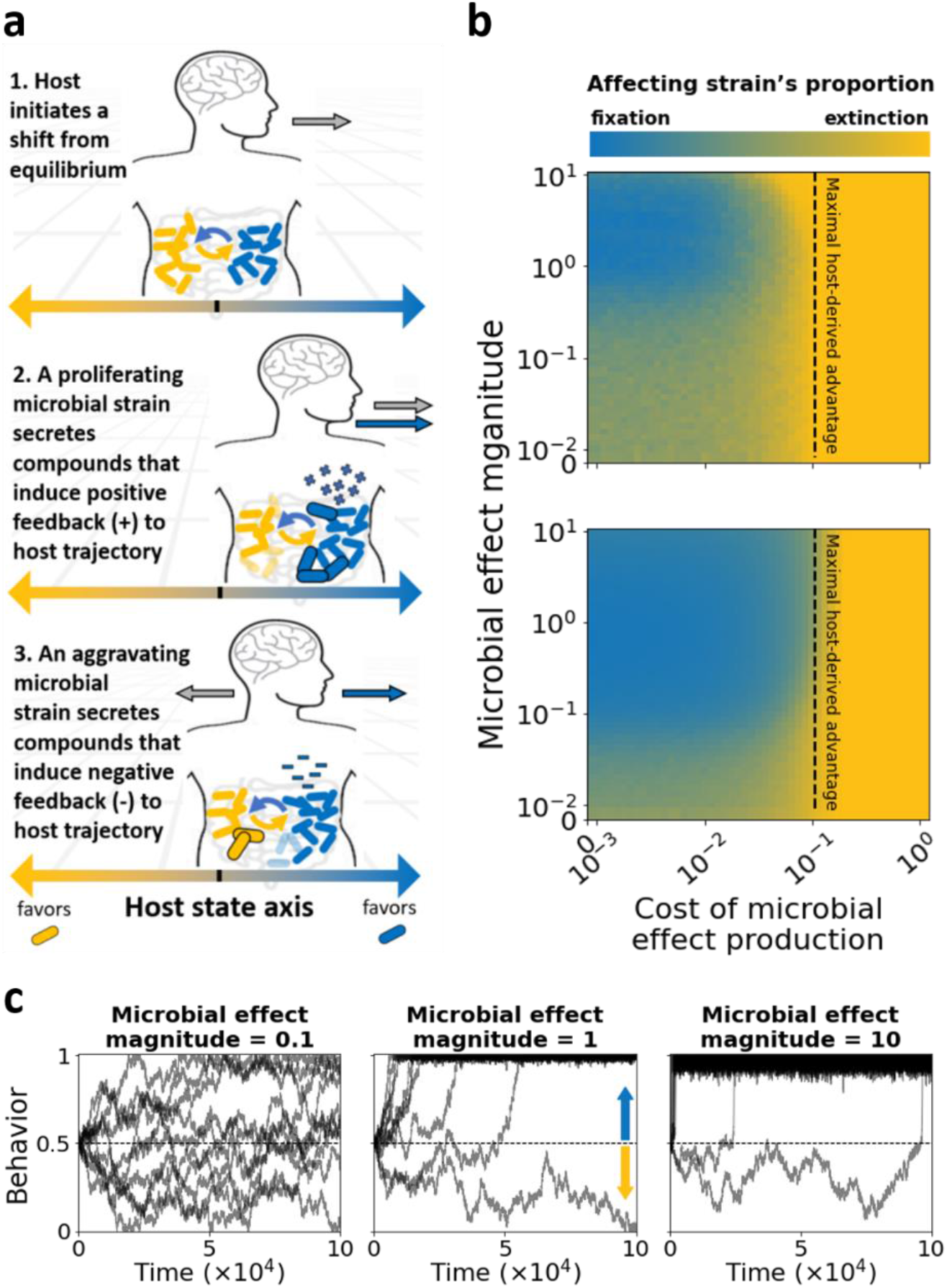
A microbe’s effect on its host’s state may be beneficial when it provides an advantage over a competitor that outweighs the cost of producing the effects. **(a)** Model Illustration. We model competition between two microbial strains for host resources, which are derived from the host behavior. Host behavior is modeled as a random walk along the [0,1] segment, starting at 0.5. The microbial strains are represented by coordinates on that same segment, while the distance between their coordinates and the behavior coordinate represents the fitness of each strain under that host behavior (see Methods). One of the two microbial strains (blue) has the potential to affect its host behavior: it secretes positive feedback when proliferating, inducing the host to continue its behavioral trend, and secretes negative feedback when declining, inducing the host to reverse its behavioral trend. These secretions are proportional to the strain’s abundance (see Methods). **(b)** Heat maps presenting the proportion of the strain that affects its host, after 100,000 time points, as a function of the effect magnitude and the cost of producing the effect. In the upper panel we set the intra-strain competition factor (*s*) to 0.01, and in the bottom panel to 0.1. The vertical dashed lines represent the maximal advantage that the affecting strain can gain from the host behavior, relative to its competitor (0.1 in these simulations); thus it is not expected to succeed when the cost is greater (see Methods). Each pixel represents the average result of 500 simulations. **(c)** Time series examples showing the host behavior as function of time over 100,000 time points. *s* = 0.1 and the cost of effect production is 0.09. In (b) and (c) the coordinates of the affecting strain and its non-affecting competitor were set to 0 and 1 respectively. The line at 0.5 is the starting position of the behavior. See Methods for a detailed description of the model.

We employed this framework to study the potential effect of microbiome behavioral feedback on host addictions. Expanding our initial model, we considered a microbiome composing *N* microbial strains, with features modeled as 2D-coordinates in a unit-sphere. We assume that the microbes compete with each other for resources from the host, and that each microbial strain has the potential to affect its host’s behavior (Fig 2a). In each simulation we randomly assign each strain its coordinate (uniformly across the unit-sphere) and its effect on the host (denoted by *d*_*i*_ for strain *i*; sampled from a distribution with mean *E*[*d*]; see Methods). We focused on the scenario of a host initiating a gradual change in behavior (*Addiction*), until it reaches some peak behavior level, where it stays for a period, followed by a gradual return towards the initial behavior (*Withdrawal*). We define the peak behavior by setting 0 < *R* < 1 as the maximal addiction severity. This simple addiction scenario serves as a baseline to which we compare the potential impact of the microbiome. As the microbial attributes are randomly chosen, we set the domain of the host behavior to be the [0,1] segment along the x-axis. This enables us to consider the host behavior-coordinate as the severity of the addiction or the degree of consumption of an addictive substance, and its effect on the microbiome.

**Figure 2.**
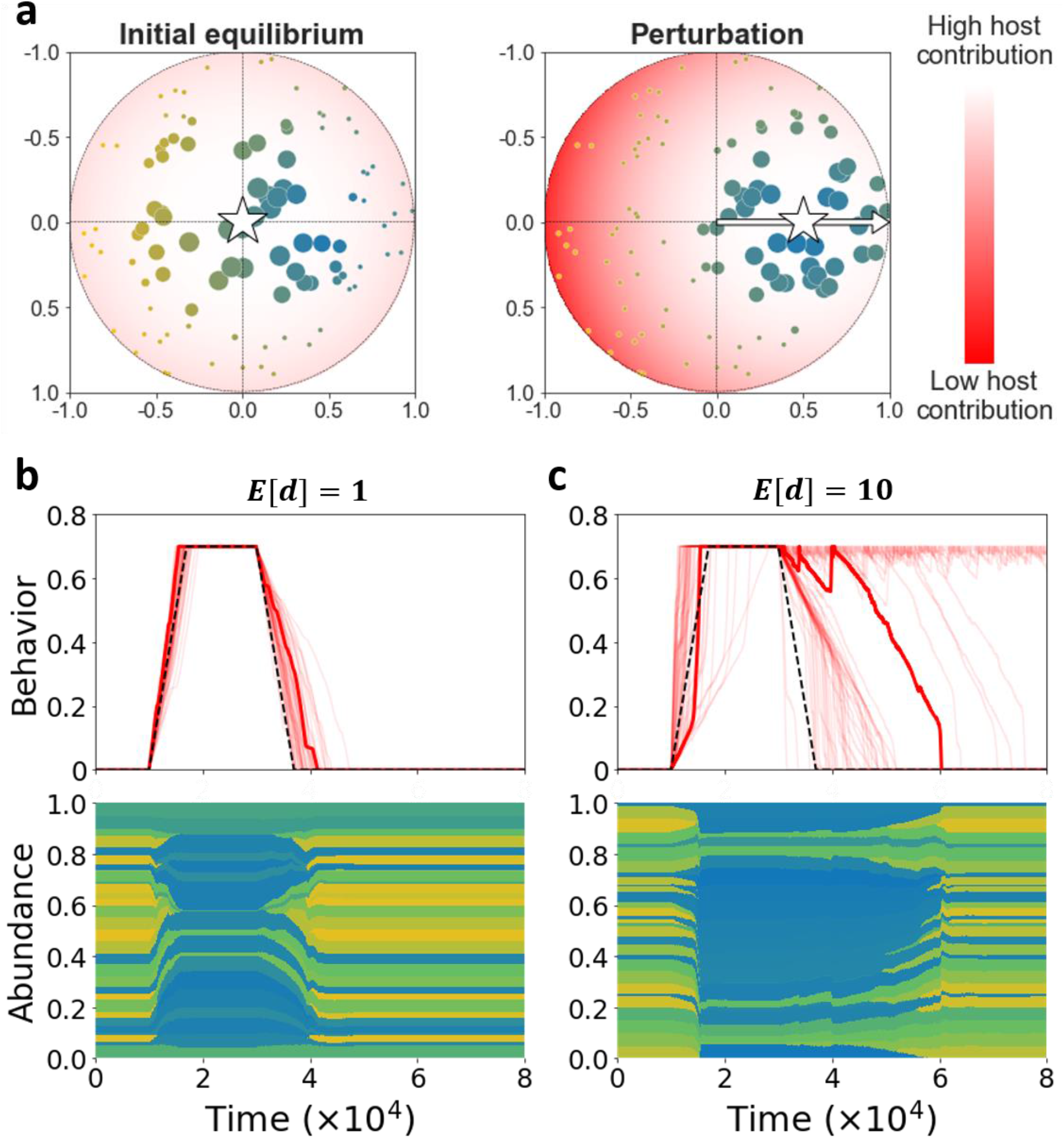
Microbiome effects on host behavior may significantly decelerate the withdrawal stage. **(a)** The host behavior is represented by a coordinate in the 2D unit sphere (star). Microbial strains (colored dots) are characterized by features, represented as coordinates in that same 2D unit sphere. The host contribution to the growth of each strain is a function of the distance between the host behavior and the strain’s features (see Methods). Thus, the microbial coordinates represent the access to host-derived resources, as function of the host behavior. The illustration demonstrates how a perturbation in the host behavior (movement of the star) produces a change in the contribution of the host to each microbial strain, which affects the microbiome composition. The color and size of each dot represents the strain’s feature-distance from the perturbed host behavior (bluer is closer) and the strain’s proportion within the microbiome, respectively. **(b, c)** Simulation examples. Upper panels show a collection of simulation results: the change in behavior over time without microbiome effects (dashed black) and with (red; 20 runs in each panel), for different mean microbiome effect magnitudes (*E*[*d*] = 1, 10). The bold red curve represents a randomly selected simulation run, for which the microbiome composition along time is plotted in the bottom panels. Each stripe (yellow-blue scale) represents a microbial strain, while the width of the stripe represents the temporal proportion of the strain within the microbiome. As in (a), the color of each stripe represents the strain’s feature-distance from the perturbed host’s behavior. The number of strains (*N*) is set to 100.

We find that intra-microbiome competition, combined with microbial effects on host behavior, can lead to either acceleration or deceleration of the addiction process, while significantly exacerbating the withdrawal phase under a wide range of conditions. We assume that before the addiction begins, the host maintains a rich and diverse microbiome. Thus, when the host initiates a change in its behavior, equal numbers of microbial strains are expected to favor or disfavor the behavioral change. However, as the change in the host behavior continues, microbial strains that are favored by the altered conditions increase in proportion, and thus their feedback effect becomes stronger, inducing the host to continue in its trajectory. Accordingly, we find that the microbiome effect on host behavior can lead both to acceleration and deceleration of the addiction, depending on the distribution of microbial features and the magnitudes of their effects. In contrast, we find that the microbiome behavioral effect decelerates the withdrawal stage in most cases. This is because during the addiction the microbiome shifts towards a less diverse community, constructed from the strains that proliferate under the new host behavior. Thus, when the withdrawal stage is initiated by the host, a significant proportion of the microbiome resists the behavioral change, decelerating the withdrawal across a very wide range of conditions (Fig. 2b,c). As in many models of complex systems, this kind of host-microbiome interaction can generate a positive feedback loop that drives the system out of balance, and in severe cases it crosses a tipping point ^33,34^.

We further investigated the effects of the microbiome richness on the addiction and withdrawal processes. We examined a wide range of numbers of microbial strains in the system, from 10 to 1,000. The characteristics of each microbial strain are represented by two parameters, chosen at random: first, its ability to affect the host behavior; second, the strain’s change in proliferation resulting from the host behavior. Hence, each strain in our model can represent an arbitrary taxonomic classification, from different phyla (∼10 in the human microbiome ^35^) to different species (up to 1,000 in the human microbiome ^36,37^), or more generally, classification into units that share specific attributes, regardless of the phylogeny. To consider not only the overall duration of these processes but also the quantitative changes in behavior, we denoted the integral of the behavior level over time for each phase by *𝜙*(*Addiction*) and *𝜙*(*Withdrawal*), (Fig. 3a), and monitored them. We found that microbial richness has an effect on the withdrawal stage: withdrawal decelerates significantly with decreasing microbiome richness, and with increasing magnitude of the microbiome effect. With respect to the addiction process, the effect of microbiome richness (*N*) depends also on the mean magnitude of the microbiome effect (*E*[*d*]; see Fig. 3c,d). In cases of very high microbial richness and strong microbial effect, the microbiome may actually hasten both addiction and withdrawal (upper rows of Fig. 3c,d). A general intuition for these effects is that when the number of available strains is high, changes are easier: any change in host behavior involves a substantial number of strains that are benefitted, and thus support and intensify the host’s behavioral alteration.

**Figure 3.**
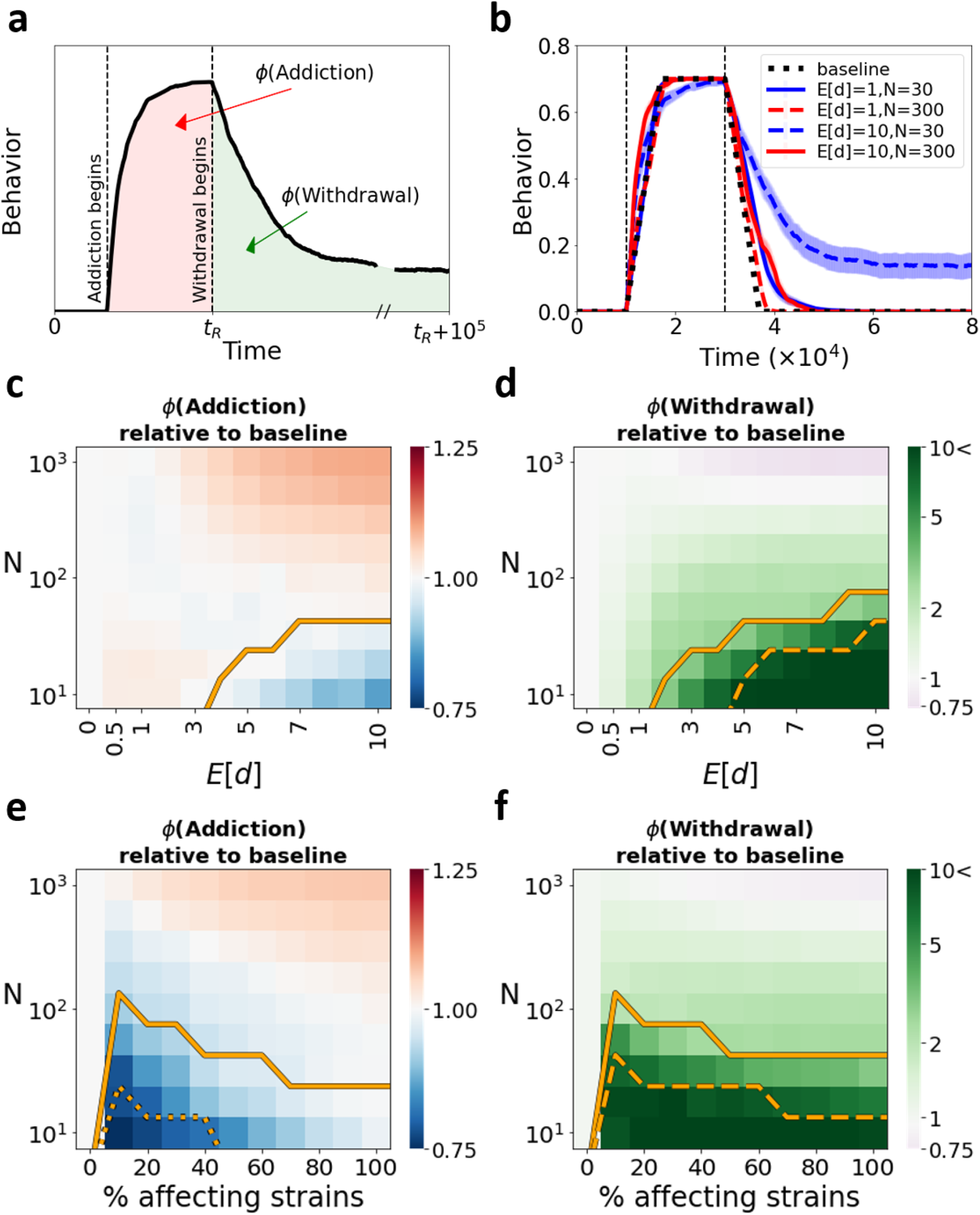
A less rich microbiome may affect the addiction process and lead to substantial aggravation of the withdrawal stage. **(a)** Illustration of a host-behavior time series. The colored markings present the integral of the behavior over time, for the addiction process (*𝜙*(Addiction); light red) and withdrawal process (*𝜙*(Withdrawal); light green). For the addiction process we consider the time from the addiction initiation until the withdrawal initiation, while for the withdrawal we considered the time from the withdrawal initiation, until the end of the simulation, 100,000 time points after withdrawal stage initiation. **(b)** Average host behavior over time is plotted for the baseline case of no microbiome effect (dashed black) and for cases that include the microbiome effect (red-blue curves), considering two mean microbial effect magnitudes (*E*[*d*]) and two numbers of available strains (*N*). Each curve presents the average of 50 simulation runs. **(c, d)** Heatmaps presenting the fold increase or decrease in *𝜙*(Addiction) (c) and *𝜙*(Withdrawal) (d) relative to the baseline case of no microbiome effect, as functions of *N* and *E*[*d*]. Each pixel in the heatmap represents the average of 1,000 simulations. **(e, f)** Heatmaps presenting the fold increase or decrease in *𝜙*(Addiction) (e) and *𝜙*(Withdrawal) (f) relative to the baseline case of no microbiome effect, as a function of *N* and of the percentage of strains that can affect the host behavior. Each pixel in the heatmaps represents the average of 1,000 simulations. To keep the mean of the overall manipulation strength of the microbiome constant and vary only the proportion of strains that affect the host behavior, we set the mean magnitude of the microbes’ effects *E*[*d*] = 5 /(*proportion of affecting strains*). Below the solid lines in (c) and (e), in more than 1% of the simulations the behavior does not reach the maximal addiction severity (*R*), and below the dashed line, more than 20% do. Below the solid line in (d) and (f) the behavior in more than 1% of the simulations does not return to the initial state at the end of the simulation, and below the dashed line, more than 20% do. *R* = *0*.*7* throughout.

We also investigated the case where only some of the strains affect host behavior by examining the impact of the distribution of microbial effects on host behavior. We first varied the proportion of strains that affect host behavior, while keeping the average of the total microbiome effect constant. Distributing the effects among fewer microbial strains resulted in deceleration of both addiction and withdrawal (Fig. 3e,f). Fixing the mean magnitude of microbial effects, then decreasing the proportion of affecting strains, resulted in a decrease in the overall microbiome effect, and accelerated the withdrawal process. Nevertheless, even when only a minority of the strains affect host behavior, the impact on addiction and withdrawal can be significant (Fig. S1). We also examined the effect of applying various levels of costs imposed by the microbial effects, and setting a constant host effect on the growth of some of the microbial strains (rather than allowing behavioral dependence): our model was robust to these changes (Fig. S2-4).

Last, we examined the effect of the microbiome’s interaction with the maximal severity of the addiction (*R;* see Fig. 2b), which represents the maximal impact of host behavior on the microbiome: higher values mean that the host can reach a state in which microbiome compositions are more distant from the initial equilibrium. We find that as the addiction becomes more severe (higher *R*), the host behavior generates an ecological regime that leads the microbiome towards a narrower niche with lower diversity. After establishment, the new microbiome composition may strongly reject any attempt to make a change, thus slowing down the withdrawal process. This dynamic increases with the magnitude of the effect of the microbiome on the host behavior, and decreases with microbiome richness. When the microbiome is richer and/or its effect on the host is relatively weak, only a substantial alteration in the microbiome composition results in a significant aggravation of the addiction (Fig. 4a). We also saw an effect of the maximal addiction severity (*R*) on the occurrences of microbiome-induced relapses. We defined a relapse as an aggravation in the addictive behavior (increase in the behavior coordinate) that occurs during the withdrawal phase (Fig. 4b). We found that stronger microbial effects, lower microbiome richness, and higher addiction severity all result in stronger and more frequent relapses (Fig. 4c).

**Figure 4.**
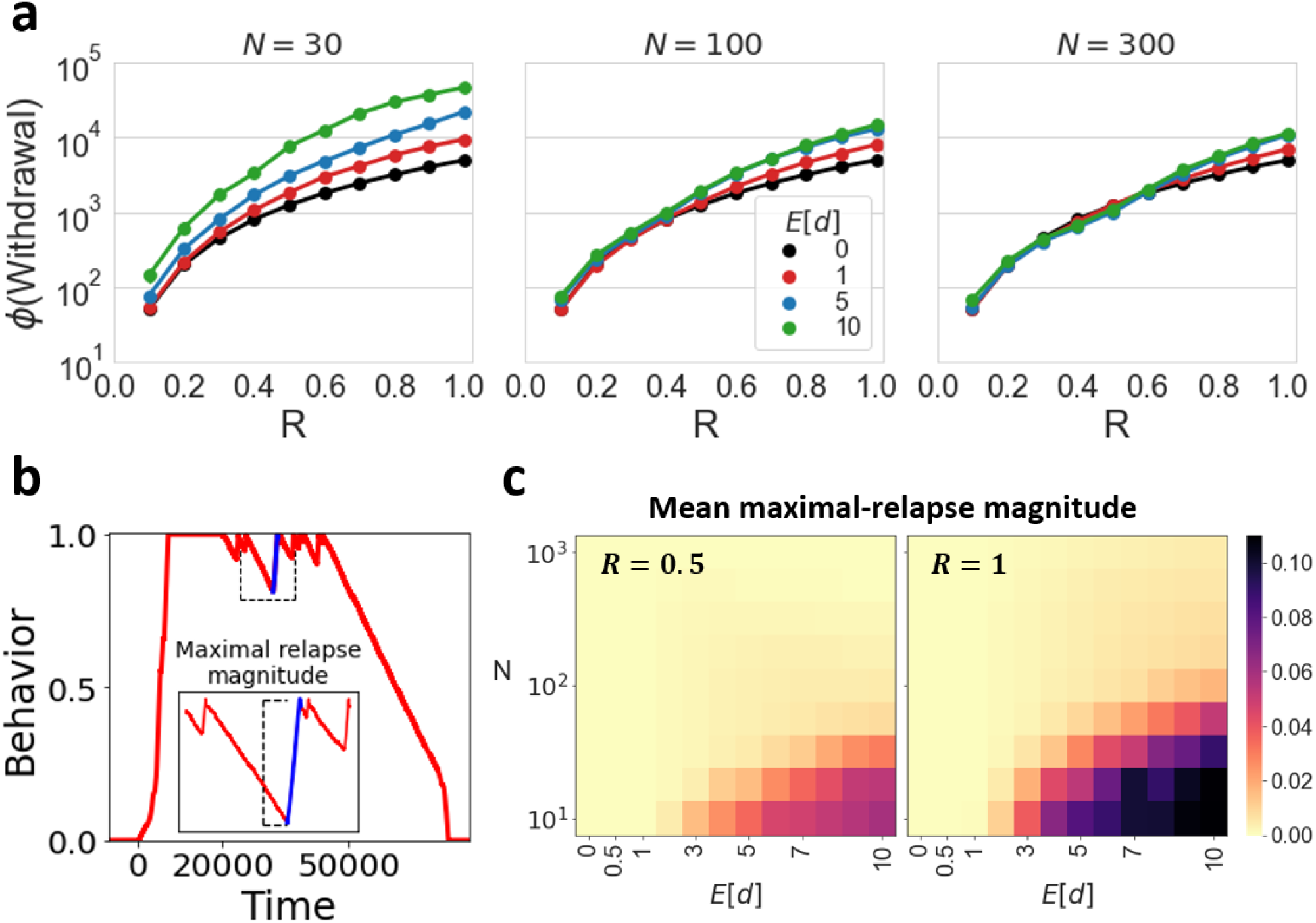
Microbiome exacerbating effect on host withdrawal increases with addiction severity. **(a)** The integral of the behavior over time during the withdrawal stage (*𝜙*(Withdrawal)) is plotted as a function of the maximal addiction severity (*R*), for several numbers of available strains (*N*) and the mean magnitudes of the effects (*E*[*d*]). Each dot represents the average of 1,000 simulations. **(b)** Relapse schematic example. We define a relapse as an increase in the addictive behavior that occurs after the withdrawal phase has begun. In each simulation we define the maximal-relapse magnitude as the maximum among the differences between all coordinates of the behavior in the withdrawal phase, and the coordinates of behavior that follow. **(c)** Heatmaps presenting the mean maximal-relapse magnitude as a function of *N* and *E*[*d*], for different maximal addiction intensities. Each pixel in the heatmaps represent the average of 1,000 simulations.

## Discussion

We hypothesized that within-microbiome competition may lead to evolution of microbial effects on host behavior, and that these effects could play a significant role in host addictive behavior. Our results demonstrate that microbiome feedbacks to host behavior can aggravate addictive behaviors, making withdrawal attempts more difficult and leading to higher risk of relapses. Microbiome richness was found to be a key parameter in the process, with low richness resulting in prolonged addictions.

Our framework includes both within-microbiome competition and host-microbiome interactions and can be extended in several ways. Other microbial interactions can be incorporated besides competition, e.g. cooperation and exploitation. Additionally, the interactions between host and microbiome can be modeled by more complex functions, e.g., threshold functions or non-monotonic functions, including rugged landscapes ^38,39^. Several different forms of microbial effects on the host behavior can be investigated, including specific reactions to certain host behaviors ^4^ (e.g., microbial reaction to exposure to opiates leading to secretion of compounds that generate host tolerance and hence induce addiction). Our model assumes a more general interaction, where the microbes can provide feedback – either positive or negative – on host behavior, according to the effect of host behavior on the microbes. Moreover, we consider a simple model of host addictive behavior, and focus on the effect of the microbiome on the process. Potential extensions may also include integrating host-brain mechanisms of addiction (reward sensitization, associative learning, conditioned reinforcers, impulse control, etc.), where the microbiome is able to affect these processes, as well as short-term and long-term effects of the addiction from both the host and microbiome perspectives.

In our model, the microbial effect on host behavior can be considered a form of public goods – microbial secreted products that benefit the strains that proliferate in the new host-mediated conditions. Numerous studies have investigated public goods secretions in microbial communities, demonstrating that such behaviors can be maintained in a population despite their direct cost to the secreting microbe, while also analyzing the conditions for the evolution of such behaviors ^40-42^.

Targeting the microbiome may be a novel direction for addiction treatment. Our model predicts that increased microbiome richness and functional diversity could contribute to addiction mitigation and prevention, and suggests that hosts with very low microbiome richness may be at greater risk for addiction and for relapse during the withdrawal period. In this context, stress and anxiety are factors that are associated with both low microbiome diversity and host addictive behaviors ^43-46^, suggesting that the interplay between host physical and mental state, microbiome composition, and addictions would be interesting to investigate. Future experiments will be required to decipher the mechanisms underlying microbiome involvement in host addictive behavior and brain function. Investigating the microbes that may benefit from the addiction may reveal potential candidates for interventions.

The framework presented in this study can be generalized and used to investigate other host-microbiome interactions with various implications on host behavior and wellbeing. For example, our framework could be relevant for studying host-microbiome interactions with respect to the host immune system, where the immune system of the host shapes the microbiome composition, and the microbiome affects the development and maintenance of the immune system ^47,48^. Moreover, this framework can be used for modeling complex ecological systems, incorporating numerous species that compete with each other, as well as environmental factors that affect the competition dynamics between these organisms, but are also altered by them ^49^. In this context, our model can be considered as a form of niche construction ^50^, where a community of species acts to construct a favorable niche in a certain environment; in our case, the environment is by itself an organism.

Better understanding of host-microbiome interactions with respect to addictive behaviors may uncover additional mechanisms behind addictions and generate novel strategies for treatment. Our results call for empirical studies – identifying key microbial strains, uncovering mechanisms of microbial effects, and testing the association between microbiome richness and functional diversity on the one hand and addictive behavior on the other.

## Methods

We model a host interacting with its microbiome, where the host behavior affects the microbiome composition and the microbiome affects the host behavior. We consider two models using the same framework. First, we examine the potential advantage of microbes’ ability to affect their host’s behavior, by modeling the growth dynamics of two microbial strains residing in a host: one strain affects its host behavior, and a second strain does not. Second, we examine the potential role of microbial effects on addiction, accounting for a microbiome community that comprises *N* microbial strains, where some or all of the strains can affect the host’s behavior.

### 1. Host baseline behavior

We represent the host baseline behavior as coordinates (denoted by 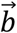) in a 2-dimensional unit-sphere, termed the microbiome-behavior-space. The host’s baseline behavior changes over time according to a pre-defined pattern. This baseline serves as a null model, representing the host behavior free of the microbiome’s effect. We later add the microbiome effect to this baseline.

#### 1.1 Two-strain competition model’s baseline

For the two-strain competition, we model the host baseline behavior as a random walk along the segment [*0,1*] on the X-axis of the microbiome-behavior-space; this can represent, for example, the consumption of a substance. Each simulation starts in the middle of that range, at 0.5. The step at each time point *t*, denoted by *𝜎*_*RW*_(*t*) is randomly drawn from a double-exponential (Laplace)^51,52^ distribution with mean *0* and scale *𝜎*. In the simulations that yielded the results for figure 1 we used *𝜎* = 10^−*3*^.

#### 1.2 Addiction model’s baseline

We focused our analysis on host baseline behavior that follows a simple addiction scenario, including three behavioral phases (see Fig. 2b):

1. **Initial equilibrium**. 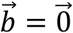 This phase ends when the host’s microbiome composition stabilizes.
2. **Addiction**. Alteration in the baseline behavior. We simulate the change in the host baseline behavior as a series of positive movements along the X-axis. At each time point *t*, a step size, denoted by *𝜎*_*A*_(*t*), is randomly drawn from an exponential distribution with mean *𝜎*. We assume that the change in the host behavior is bounded by distance *R* from the origin, representing the maximal addiction severity. Thus, the host-behavior coordinates shift gradually from (*0,0*) up to (*R, 0*), and stay at (R,0) until the end of this phase, *𝜏* time steps after its initiation.
3. **Withdrawal**. The host reverses its behavioral pattern, by moves of (−*𝜎*_*A*_(*t*)) on the X-axis, with *𝜎*_*A*_(*t*) randomly drawn from exponential distribution with mean *𝜎*. The change in behavior is bounded below by 0. This phase ends when the microbiome composition and host behavior stabilize, or after 100,000 time steps from the initiation of the phase.
4. In the simulations that yielded the results for the figures (2-4) and the supplementary figures we used *𝜎* = 10^−*4*^ and *𝜏* = 20,000.

### 2. Microbiome composition

We model a population of *N* microbial strains inhabiting a host. Each strain has its unique features, modeled as coordinates in the microbiome-behavior-space. We denote the vector of these coordinates for each strain *i* by 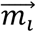. The following system of ordinary differential equations describes the change in the frequency of strain *i*, along time^26,53^:

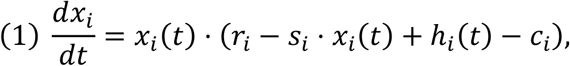

where *x*_*i*_ denotes the proportion of strain *i* within the microbiome; *r*_*i*_ denotes the intrinsic baseline growth rate of strain *i*; *s*_*i*_ denotes intra-strain competition; *h*_*i*_ denotes the effect of the host behavior on the growth of strain *i*; *c*_*i*_ denotes the constant cost of feedback production experienced by strain *i*. We used the forward Euler method to simulate the dynamics of this system of equations, with time steps of size 1. At each step of the simulation, we normalized the *x*_*i*_ values by their sum, to give for the frequency of each strain.

We also assume a constant inflow of all the *N* possible microbial strains at rate *µ*/*N* for each strain, thus our results do not rely on extinction and permanent disappearance of strains from the system (we used *µ* = 10^−*8*^ throughout). At each time step, the strain proportions are calculated using (1), followed by inflow and normalization.

In the simulations that yielded the results for figures 2-4 and the supplementary figures we used *r* = 0.1, *s* = *1*. For figure 1 we used *r* = 0.1 and *s* = 0.1,0.01. We analyze the effect of *c*_*i*_ *≠ 0* in figure 1 and in figures S2, S3, and set *c*_*i*_ = *0* in the simulations that yielded the rest of the results.

#### 2.1 Two-strain competition models

For the two-strain competition model (*N* = *2*) the features of each strain are manually defined and specified in the results. One strain affects the host behavior, and it thus pays the cost of feedback production, while the other strain does not affect the host behavior and does not pay the cost.

#### 2.2 Addiction model

For the addiction model the features of each strain are drawn randomly within the microbiome-behavior-space (from a uniform distribution on the 2D unit sphere), at the beginning of each simulation. We consider a microbiome community that comprises *N* microbial strains, where some or all of the strains can affect the host behavior.

### 3. Host-behavior effect on the microbiome composition

We focus on aspects of host behavior that regulate the ecology of the microbiome and thus affect the microbiome composition. The host-derived resources obtained by each strain are determined by a monotonically decreasing function of the Euclidean distance between the host and the microbial strain, in the microbiome-behavior-space. In this context we focus on the microbial features relevant for the host-microbe interaction. We thus define *h*_*i*_ at time *T* as follows:

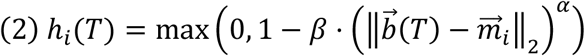

In the simulations that yielded the results for figures (1-4) and for the supplementary figures we used *α* = *3, β* = 0.1. Fig. S6 demonstrates the robustness of our results to these parameters. We also examined the effect of setting a constant host effect on the growth of some microbial strains (rather than having the effect be behavior-dependent), and our model was robust to this change (Fig. S4).

### 4. Microbiome effect on host behavior

We assume that all or some of the microbial strains can affect their host’s behavior, and that the microbes react to positive and negative changes in their environment by producing and secreting compounds that are perceived by the host as positive or negative feedback (e.g. reward or aversion). We denote by 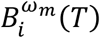 the slope of the linear regression on the proportions of strain *i* in the past *ω*_*m*_ time points. Namely, 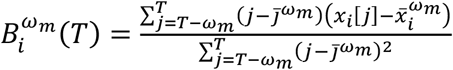, where 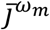 average of the time-point indicators (*T* − *w*_*m*_,*T* − *w*_*m*_ + 1, …, *T*), *x*_*i*_[*j*] is the proportion of strain *i* at time point *j*, and 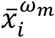 is the average of strain’s *i*’s proportions during the time period *T* − *ω*_*m*_ through *T*. We then denote by 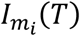 the condition of microbial strain *i* at time *T*, defined as follows:

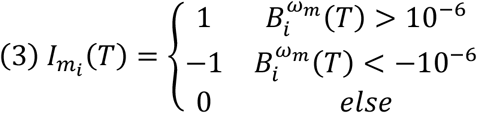

Here, the condition of strain *i* is positive, negative, or neutral (and thus it secretes positive feedback, negative feedback, or no feedback at all) depending on the slope of the linear trajectory of its proportion in the past *ω*_*m*_time points. Slopes that are very close to zero (between −10^−*6*^ and 10^−*6*^) are considered neutral in their effect, representing accuracy limitations in evaluation. These feedbacks affect the host behavioral change later on.

We denote by 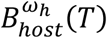 the slope of the linear regression on the host behavior in the past *w* time points (considering its moves along the X-axis; similarly to the calculation in the previous paragraph, only that the regression is of the host-behavior coordinate along the X-axis). We then denote by *I*_*b*_(*T*) the behavioral trajectory of the host at time *T*, defined as follows:

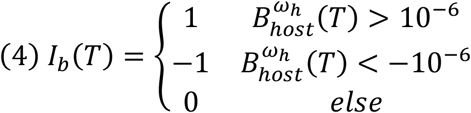

Hence, the host trajectory – away from the center, towards the center or neutral – depends on the slope of its linear trajectory over the past *w*_*h*_ time points.

In the simulations that yielded figures (1-4) and for the supplementary figures we used *w*_*h*_ = *w*_*m*_ = 10. See Fig. S5 which demonstrates the robustness of our results to these parameters.

Combining the microbial secretions and the behavior trajectory, the direction of microbe *i*’s effect on the host behavior at time *T* can be described by the sign of the product 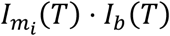. This term is positive when strain *i* influences the host to advance on the addiction path (e.g., to increase doses of the consumed substance), in one of two scenarios: strain *i* providing positive feedback 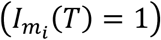 to host advancement on the addiction path (*I*_*b*_(*T*) = *1*), or strain *i* providing negative feedback 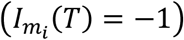 to host withdrawal (*I*_*b*_(*T*) = −*1*). The term 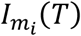 is negative when strain *i* influences the host to take a step backwards on the addiction path (e.g., to reduce consumption of the substance); and when it is zero, strain *i* does not affect host behavior.

The total effect-strength of each strain, at time *t*, is that strain’s proportion within the microbiome, multiplied by(|*𝜎*_*A*_(*t*)| ⋅ *d*_*i*_), where *d*_*i*_ denotes the magnitude of the effect of strain *i* on the host, and |*𝜎*_*A*_(*t*)| is the baseline behavioral step size at time *t*. The microbial effect magnitudes (*d*_*i*_, *i ∈* {1, *…, N*}) are chosen at random from an exponential distribution with mean *E*[*d*] at the beginning of each simulation, and *𝜎*_*A*_(*t*) are drawn at each time step, as explained in section 1.2.

The effect of the entire microbiome at time *T*, denoted by *M*(*T*), is the sum of all strains’ effects:

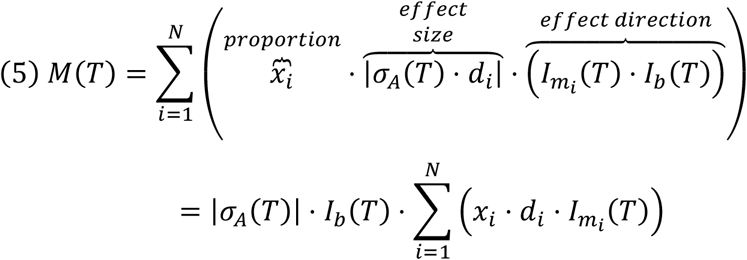

The change in host behavior from time *T* to time *T* + 1 is defined by the sum of the host baseline behavior step and the microbiome-induced step. We denote by *b*_*1*_ the first coordinate of the host behavior (corresponding to the X-axis, along which the host moves) and define the host behavior at time *T* + 1 as follows:

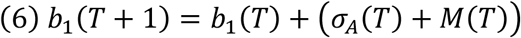

## Acknowledgements

We wish to thank Mark Tanaka and David Zeevi for insightful discussions and comments on the manuscript.

## Authors’ contributions

O.L.-E., L.H., D.K., Y.J. and M.W.F designed the study. O.L.-E. and L.H. formulated the model. O.L.-E. wrote the simulation code and conducted the analysis under the supervision of L.H.. O.L.-E., L.H., Y.J, D.K, and M. W. F. analyzed the results and wrote the manuscript.

