## Supplementary figures for "Addictions may be driven by competition-induced microbiome dysbiosis"

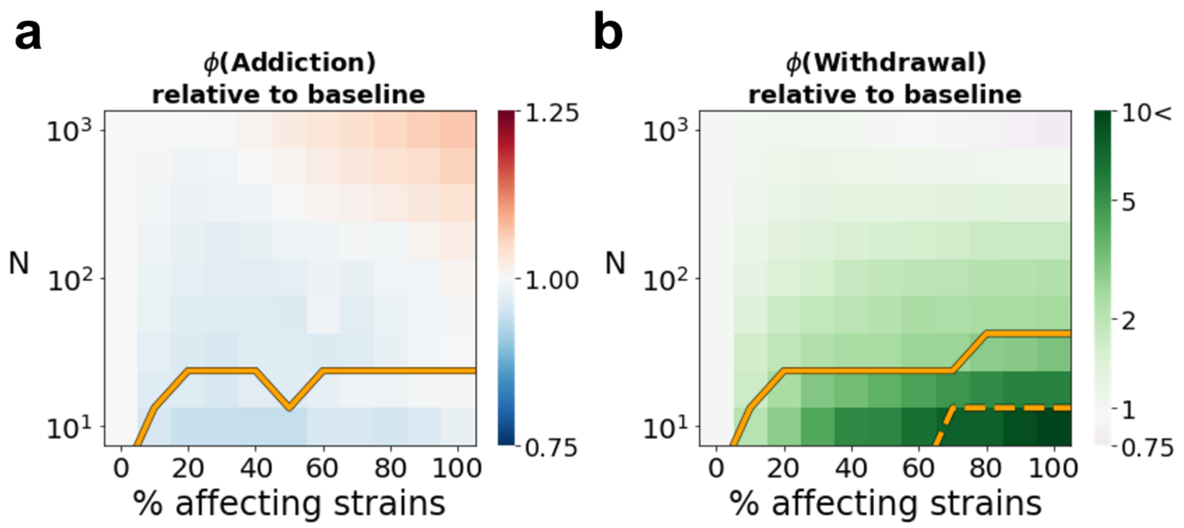


**Figure S1. The microbiome can strongly affect the addictive behavior, even when the majority of the strains cannot affect the host behavior.** Heatmaps presenting the fold increase or decrease in $\phi$(Addiction) (a) and $\phi$(Withdrawal) (b) relative to the baseline case of no microbiome effect, as function of $N$ and of the percentage of strains that can affect the host behavior. Each pixel in the heatmaps represents the average of 1,000 simulations. The figure is similar to Fig. 3e,f in the main text, except for the mean effect magnitude of the affecting microbes ($E[d]$) which in the simulations presented in the current figure was set to 5, throughout. Below the solid lines in (a) more than 1% of the simulations do not reach the maximal addiction severity ($R$). Below the solid line in (b) the behavior in more than 1% of the simulations does not return to the initial state at the end of the simulation, and below the dashed line, more than 20%. $R = 0.7$.

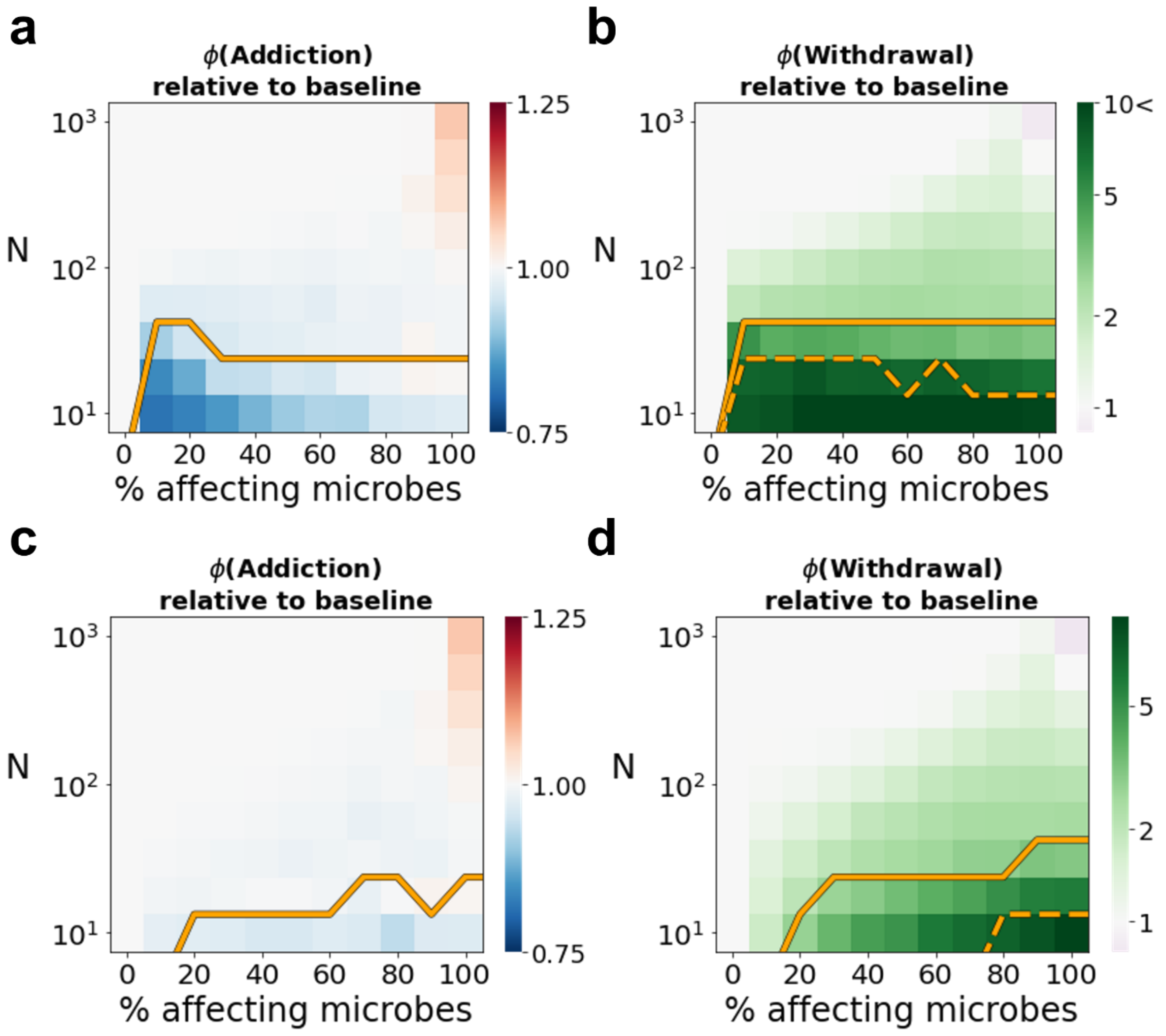


**Figure S2. The microbiome can strongly affect the addictive behavior, even when only part of the microbiome affects the host behavior, and including when the microbial production of the effect is costly.** Heatmaps presenting the fold increase or decrease in $\phi$(Addiction) (a,c) and $\phi$(Withdrawal) (b,d) relative to the baseline case of no microbiome effect, as function of $N$ and of the percentage of strains that can affect the host behavior. Each pixel in the heatmaps represents the average of 1,000 simulations. The figure is similar to Fig. 3e,f in the main text, but the simulations presented in the current figure included cost of 0.03 conferred by the microbes that had the ability to affect host behavior (see Methods, and equation 1). For the simulations that generated panels (a) and (b) we set $E[d] = 5\ /(proportion\ of\ affecting\ strains)$, while for panels (c) and (d) we set $E[d] = 5$. Below the solid lines in (a,c) more than 1% of the simulations do not reach the maximal addiction severity $(R)$. Below the solid line in (b,d) the behavior in more than 1% of the simulations does not return to the initial state at the end of the simulation, and below the dashed line, more than 20%. $R = 0.7$.

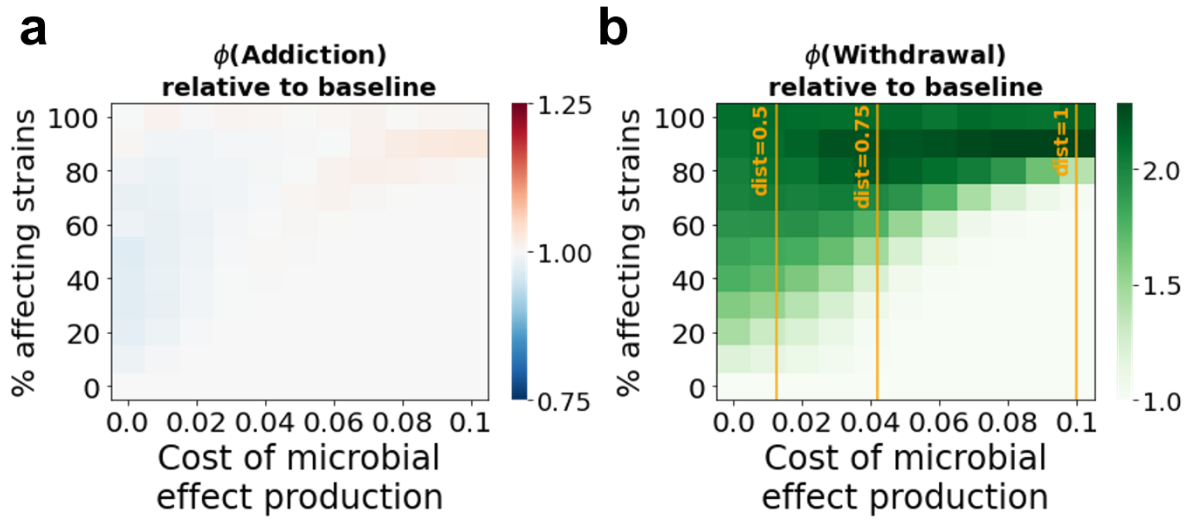


**Figure S3. Microbiome effect on host withdrawal can be significant even when only part of the strains can produce the effect, and even when this production is costly.** Heatmaps presenting the fold increase or decrease in $\phi$(Addiction) (a) and $\phi$(Withdrawal) (b) relative to the baseline case of no microbiome effect. Each pixel in the heatmaps represents the average of 1,000 simulations. The vertical lines in (b) indicate the distance advantage (on the microbiome-behavior-space) that an affecting microbe must have in an optimal host behavior, relative to non-affecting strain, in order to compensate for the cost of microbial effect production. $R = 0.7, N = 100, E[d] = 5$.

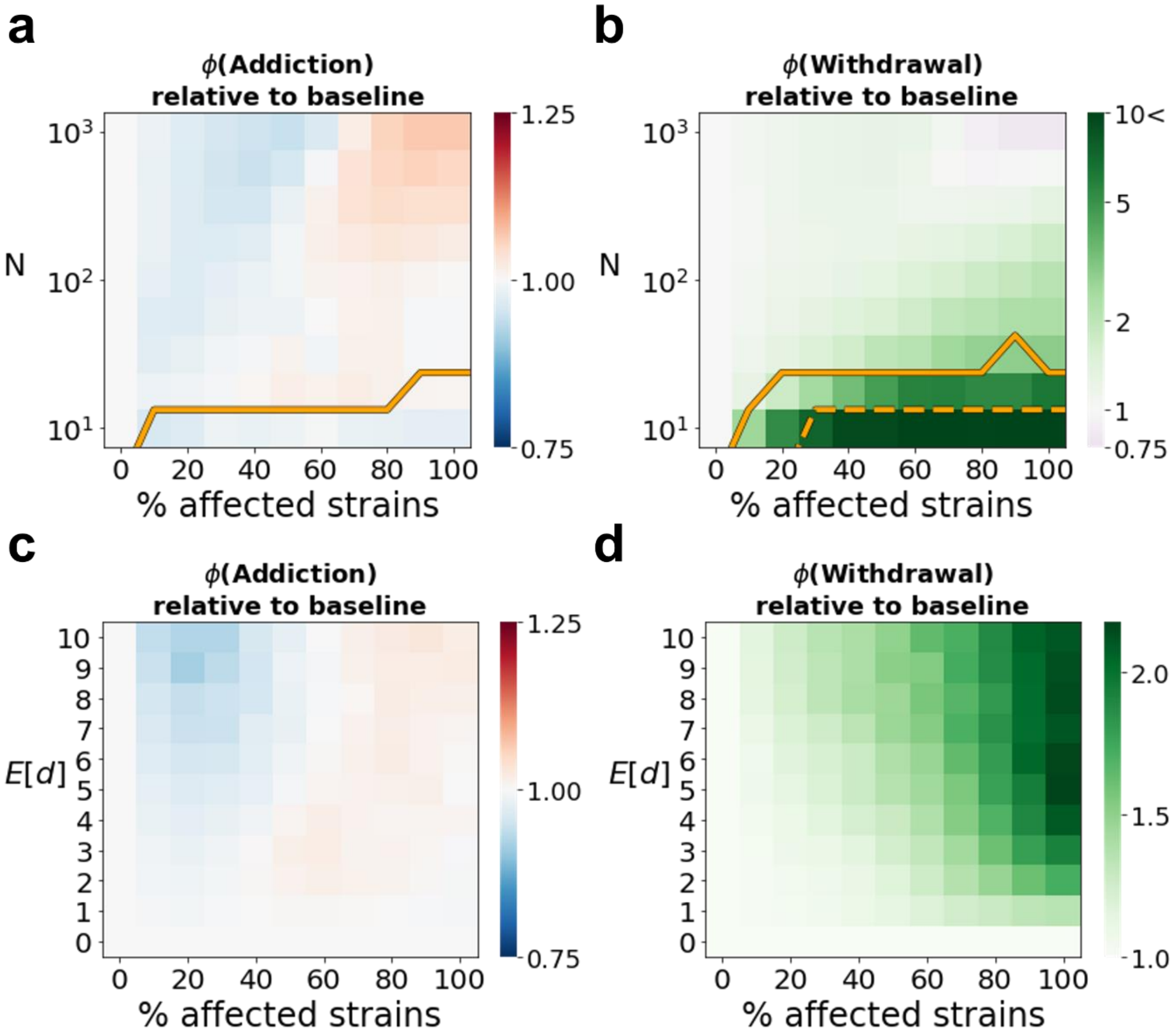


**Figure S4. Microbiome effect on host addiction and withdrawal may be significant even when only part of the microbiome is affected by the changes in the host behavior.** Heatmaps presenting the fold increase or decrease in $\phi$(Addiction) (a,c) and $\phi$(Withdrawal) (b,d) relative to the baseline case of no microbiome effect, as function of the proportion of strains that are affected by the host changes in the behavior (x-axis), $N$ (on the y-axis of panels a,b) and $E[d]$ (on the y-axis of panels c,d). In the simulations executed for these results, the host factors that affect the strains' growth ($h_i$; see equation 1 in the Methods) where set to remain constant throughout the simulation. These strains (following the proportions as indicated in the x-axis) were randomly chosen at the beginning of each simulation and their $h_i$ value for the entire simulation were determined according to the distance between each strain feature-coordinates, and the initial behavior coordinate (set to the origin). Each pixel in the heatmaps represents the average of 1,000 simulations. Below the solid line in (a) more than 1% of the simulations do not reach the maximal addiction severity ($R$). Below the solid line in (b) the behavior in more than 1% of the simulations does not return to the initial state at the end of the simulation, and below the dashed line, more than 20%. $R = 0.7$. In panels (a) and (b) $E[d] = 5$ while in panels (c) and (d) $N = 100$.

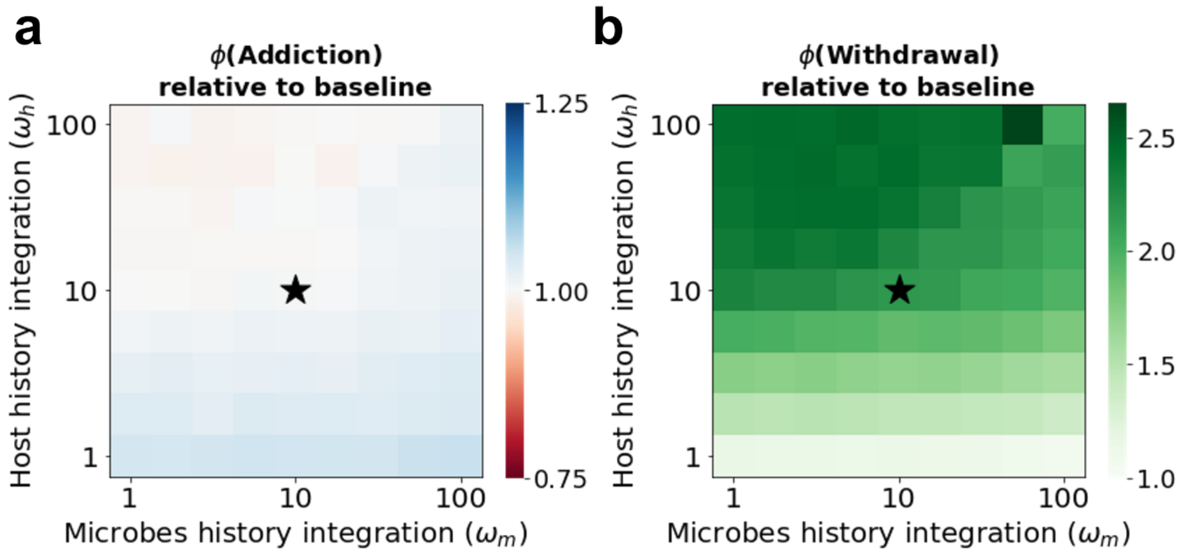


**Figure S5. Robustness of the results to the selection of the length of history time frame considered in the host and microbiome determination of trajectory.** Heatmaps presenting the fold increase or decrease in $\phi$(Addiction) (a) and $\phi$(Withdrawal) (b) relative to the baseline case of no microbiome effect, as function of $\omega_h$ and $\omega_m$ (see Methods). Each pixel in the heatmaps represents the average of 1,000 simulations. Marked by star are the parameters that were used throughout the manuscript. $R = 0.7, N = 100, E[d] = 5$.

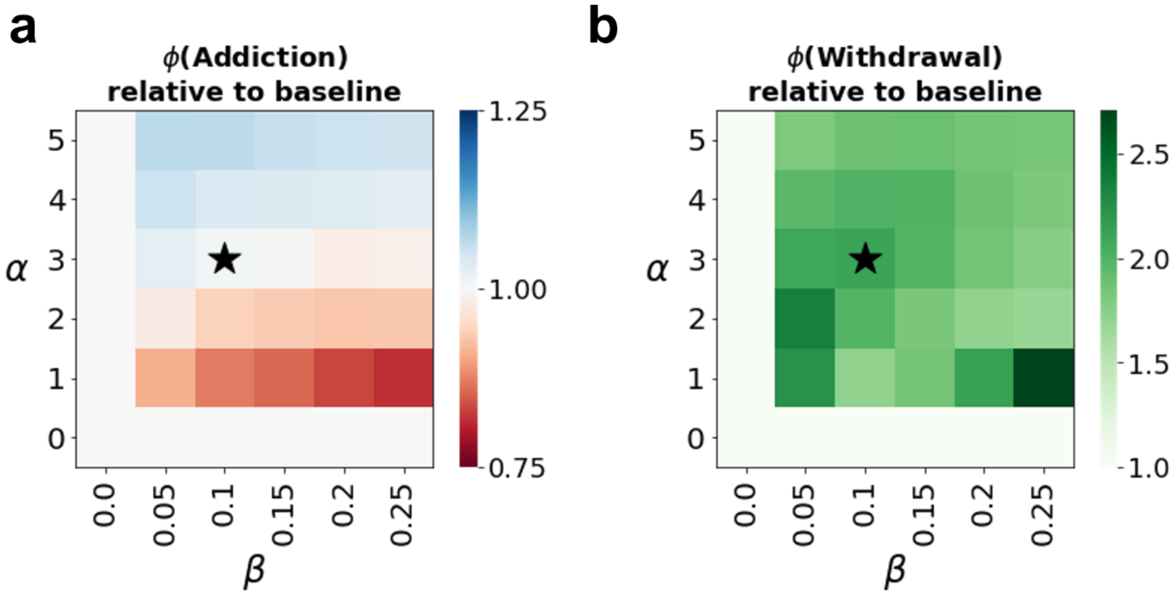


**Figure S6. Robustness of the results to the selection of the parameters controlling the host-behavior impact on the microbes' growth.** Heatmaps presenting the fold increase or decrease in $\phi$(Addiction) (a) and $\phi$(Withdrawal) (b) relative to the baseline case of no microbiome effect, as function of $\alpha$ and $\beta$ (see Methods). Each pixel in the heatmaps represents the average of 1,000 simulations. Marked by star are the parameters that were used throughout the manuscript. $R = 0.7, N = 100, E[d] = 5$.
